# Time at Home for Whom? Unequal Returns of Paid Family Leave for Infant White Matter and Language Development

**DOI:** 10.64898/2026.10.08.757668

**Authors:** Molly F. Lazarus, Charles A. Nelson

## Abstract

Paid family leave (PFL) may give parents more time with infants, ease financial strain, and support stable routines. Although PFL has been linked to child development, little is known about neural pathways that might underpin its benefits or whether associations vary across families. Using data from 274 infants in the HEALthy Brain and Child Development Study, we examined associations of maternal and paternal PFL with language ability at 12 months and white matter microstructure at 6 months. We tested whether associations varied by the corresponding parent’s education. Neither maternal nor paternal PFL was directly associated with language scores or fractional anisotropy in the left arcuate, inferior longitudinal, or uncinate fasciculus. Maternal education moderated associations between maternal PFL and fractional anisotropy in the arcuate (*p* = .014) and inferior longitudinal fasciculus (*p* = .003; *n* = 148). In both tracts, PFL was associated with higher fractional anisotropy only among infants whose mothers had relatively high education. No comparable pattern emerged for language, the uncinate fasciculus, or paternal PFL. Findings suggest that the neurodevelopmental correlates of PFL may depend on family resources rather than operate uniformly across families, underscoring the need to pair paid leave with additional supports and resources.

**Lay Summary:** Paid family leave may give parents more time at home, steadier daily routines, and less financial worry in the months after a birth, all of which could change the care an infant receives. But these benefits may not be equal across families. In a national study of infants, we examined whether paid leave taken by mothers, fathers or partners was related to early language skills and to the organization of brain pathways involved in language. Paid leave was not directly related to language scores or brain development overall. However, maternal paid leave was associated with more mature organization in two language-related brain pathways only among infants whose mothers had higher levels of education. No similar pattern was found for father or partner leave. These findings suggest that paid leave alone may not reduce early developmental disparities, and that policies offering paid leave may need to be paired with additional support for families with fewer resources.

## Paid Family Leave as an Early Developmental Context

The United States is the only high-income country without a federal policy guaranteeing paid family leave (PFL) to new parents (Van Niel et al., 2020). As of March 2023, only 27% of private-industry workers had access to employer-provided PFL (U.S. Bureau of Labor Statistics, 2023). Many families must therefore rely on unpaid, job-protected leave through the Family and Medical Leave Act or return to work before the 12 weeks recommended by the American College of Obstetricians and Gynecologists for postpartum recovery (American College of Obstetricians and Gynecologists [ACOG], 2018). Access to PFL is also strongly patterned by socioeconomic status. Workers with lower incomes and less formal education are less likely to have paid leave available, even though they may stand to benefit from it the most (Kozak et al., 2021; Van Niel et al., 2020).

A growing literature links PFL with favorable maternal and infant health outcomes. Paid leave has been associated with lower rates of maternal and infant rehospitalization, reduced postpartum depression, and greater employment stability following childbirth (Jou et al., 2018; Mandal, 2018; Rossin-Slater et al., 2013). Mothers who take unpaid or no leave are also more likely to discontinue breastfeeding earlier and to miss infant immunizations (Berger & Waldfogel, 2004). These findings suggest that PFL may shape the broader developmental context into which an infant is born. By providing protected time at home and reducing financial and employment-related strain, PFL may support caregivers’ capacity to establish consistent, responsive routines during a period of rapid infant development and increased neuroplasticity.

## Paid Leave and Early Language Development

Language may be especially sensitive to the caregiving opportunities afforded by PFL. Early language development depends on repeated exposure to speech, reciprocal social interaction, shared attention, and caregivers’ timely responses to infant vocalizations and communicative cues (Kapengut & Noble, 2020). These experiences begin well before infants produce their first words and help establish the foundations for later receptive and expressive language (Noble et al., 2015). Because paid leave may increase the amount of time caregivers are available for everyday interaction while reducing competing work and financial demands, it may create more opportunities for the types of contingent, language-rich exchanges that support early communication.

Emerging evidence is consistent with this possibility. Kozak et al. (2021) found that toddlers whose mothers took paid leave had higher parent-reported language scores than toddlers whose mothers took unpaid leave. Associations with socioemotional outcomes were also strongest among families with lower incomes, suggesting that the developmental relevance of PFL may differ across socioeconomic contexts. In a sociodemographically diverse New York City sample, Brito et al. (2022) found that infants whose mothers had paid leave showed a more mature pattern of resting electroencephalography activity at 3 months than infants whose mothers had unpaid leave, even after accounting for family income and parent-child interaction quality. Together, these studies suggest that PFL may be associated with both early language development and the neural systems that support emerging cognitive abilities. However, little is known about the neurodevelopmental pathways through which these associations may arise.

## White Matter as a Potential Neurodevelopmental Pathway

Infancy is a period of rapid white matter development. White matter pathways form the structural connections that allow information to be transmitted across distributed brain regions, supporting the emergence of increasingly complex cognitive and behavioral functions (Gilmore et al., 2018). Although white matter development is strongly influenced by biological maturation, it is also shaped by variation in early experience. Prior work has linked differences in caregiving environments and socioeconomic conditions with individual variation in white matter microstructure during childhood (Bick et al., 2015; Carozza et al., 2025).

PFL is a distal, family-level exposure, but it may influence several aspects of infants’ everyday experience that are relevant to brain development. Paid leave may affect how much time caregivers spend with their infants, the regularity of caregiving routines, the amount of social and linguistic input infants receive, and the level of stress within the caregiving environment. These experiences accumulate across infancy and may be associated with the organization of neural pathways supporting early learning. Examining white matter alongside language outcomes therefore offers an opportunity to test whether PFL is associated not only with later behavior, but also with an earlier aspect of the neural architecture that supports that behavior.

Three left-hemisphere association tracts may be particularly relevant. The arcuate fasciculus connects frontal and temporoparietal language regions and is among the most consistently implicated pathways in phonological processing and language development from infancy through childhood (Yeatman et al., 2011; Zuk et al., 2021). The inferior longitudinal fasciculus connects occipital and anterior temporal regions and supports visual-semantic integration, object recognition, and the mapping of visual information onto lexical and semantic representations (Herbet et al., 2018). The uncinate fasciculus connects anterior temporal and orbitofrontal regions and has been implicated in semantic association and lexical retrieval, in addition to its broader roles in memory and socioemotional processing (Papagno, 2011; Von Der Heide et al., 2013). Together, these pathways support complementary aspects of the developing language system.

Fractional anisotropy (FA), a diffusion MRI measure reflecting the directional organization of water diffusion within white matter, increases rapidly across infancy as fiber pathways become more organized. Examining FA in these tracts at approximately 6 months may therefore provide insight into whether PFL is associated with early variation in the structural systems that later support language.

## Socioeconomic Context

The association between PFL and infant development may not be uniform across families. Socioeconomic differences in cognitive and brain development are evident within the first year of life and widen across early childhood (Noble et al., 2015). At the same time, access to PFL is itself socioeconomically patterned. Families with fewer resources may experience greater financial strain when leave is unpaid and may therefore receive a larger developmental benefit from paid leave. Existing findings are mixed. Kozak et al. (2021) found that associations between maternal PFL and toddler socioemotional outcomes were strongest among lower-income families, whereas Brito et al. (2022) found no evidence that family income moderated associations between PFL and infant EEG. Whether maternal education, an important indicator of socioeconomic resources and of the broader learning environment available to children, moderates associations between PFL and infant white matter or language development remains unknown.

## Paternal and Partner Leave

Most PFL research has focused on mothers. However, fathers and other partners may also have access to leave through employer policies, state programs, or broader parental-leave benefits. Fathers’ leave-taking has been associated with greater subsequent involvement in caregiving, which may in turn support child development (Huerta et al., 2013). Paternal involvement has also been linked to infant neurodevelopment within the first months of life, although this literature remains limited relative to research on maternal caregiving (Kim et al., 2016).

Paternal leave may influence infant development through several pathways. It may increase the partner’s direct involvement in caregiving, provide the infant with additional opportunities for social and linguistic interaction, reduce caregiving demands on the mother, and support more stable family routines. To our knowledge, no study has examined whether paternal or partner PFL is associated with infant white matter development. Whether paternal or partner PFL is associated with early language development also remains largely untested.

## Present Study

The present study examined these questions using data from the HEALthy Brain and Child Development (HBCD) Study, a National Institutes of Health-funded, multisite longitudinal cohort following families from the prenatal period through early childhood (Nelson et al., 2024). We tested parallel associations for maternal PFL and paternal or partner PFL with two sets of infant outcomes: Bayley Scales of Infant and Toddler Development, Fourth Edition Language Standard Scores at approximately 12 months of age, and FA in the left arcuate fasciculus, inferior longitudinal fasciculus, and uncinate fasciculus at approximately 6 months of age. In pairing a behavioral outcome with an earlier structural one, our aim was not only to ask whether paid leave relates to infant development, but to begin to identify the neural mechanisms that might underpin the behavioral advantages reported in prior work.

We hypothesized that maternal and paternal or partner PFL would each be associated with higher language scores and with higher FA in the three language-relevant tracts. We also examined whether these associations varied as a function of the corresponding parent’s education, hypothesizing that families with fewer educational resources would benefit most from PFL.

## Method

### Participants

Data were drawn from the HBCD Study, an ongoing, National Institutes of Health HEAL Initiative-funded consortium study following child health and brain development across 27 recruitment sites in the United States, beginning prenatally (Nelson et al., 2024). HBCD schedules assessments within age-based visit windows. The visits used in this analysis were prenatal (V01), birth to 1 month (V02), 3–9 months (V03), 9–15 months (V04), and 10–17 months (V05).

The present data were pulled on 6/22/2026 from HBCD’s second data release and comprised 3,513 participants with demographic data collected at the prenatal study visit (V01); because infants are not yet born at V01, all childbirth characteristics were drawn from the subsequent V02 visit. Of these, 292 caregivers provided a retrospective report of maternal leave-taking at a later visit (V05). The primary analytic sample comprised the 274 infants with complete data on maternal PFL status and on the a priori covariates (gestational age at delivery, child sex, maternal education, and household income). Within this sample, 231 infants had a valid Bayley-4 Language Standard Score obtained at approximately 12 months of adjusted age (V04), and 148 infants had diffusion MRI data obtained at approximately 6 months of adjusted age (V03). A parallel sample restricted to families with complete data on paternal or partner PFL and paternal or partner education comprised 193 infants for language outcomes and 120 infants for FA outcomes.

The primary analytic sample (*n* = 274) was 51% male and identified as 61% White, 21% Black or African American, 12% two or more races, 3% Asian, and less than 1% each American Indian or Alaska Native or another race (race data missing for < 1% of the sample). Consistent with known patterns of unequal access to PFL, families in which the mother took paid leave differed from families in which she did not on several sociodemographic indicators: mothers who took paid leave had more years of education (M = 18.5, SD = 1.8) than mothers who did not (M = 16.0, SD = 2.5; p < .001), and household income was skewed toward higher brackets among paid-leave families (e.g., 42% in the $100,000–$199,999 bracket and 30% in the $200,000-or-more bracket, versus 14% and 7% respectively among families without paid leave; p < .001). Groups did not differ in gestational age at delivery (*p* = .9) or child sex (*p* = .5; Table 1).

**Table 1.** Sample Characteristics by Maternal Paid Family Leave (PFL) Status.

| Characteristic | Overall (N =<br>274)<br>M(SD) or n<br>(%) | No Paid<br>Leave (n =<br>83)<br>M(SD) or<br>n (%) | Paid Leave<br>(n = 191)<br>M(SD) or<br>n (%) | *p* |
| --- | --- | --- | --- | --- |
| **Female Sex (%)** | 133 (49%) | 43 (52%) | 90 (47%) | .5 |
| **Child race (%)** |  |  |  | < .001 |
| White | 166 (61%) | 37 (45%) | 129 (68%) |  |
| Black or African American | 56 (21%) | 32 (39%) | 24 (13%) |  |
| American Indian or Alaska Native | 2 (0.7%) | 1 (1.2%) | 1 (0.5%) |  |
| Asian | 8 (2.9%) | 1 (1.2%) | 7 (3.7%) |  |
| Two or more races | 32 (12%) | 10 (12%) | 22 (12%) |  |
| Other race | 9 (3.3%) | 2 (2.4%) | 7 (3.7%) |  |
| Missing | 1 | 0 | 1 |  |
| **Gestational age at delivery, weeks** | 38.6 (1.8) | 38.7 (1.6) | 38.5 (1.9) | .9 |
| **Maternal education, years** | 17.7 (2.3) | 16.0 (2.5) | 18.5 (1.8) | < .001 |
| **Paternal education, years** | 17.4 (2.4) | 15.9 (2.7) | 17.9 (2.1) | < .001 |
| Missing | 27 | 19 | 8 |  |
| **Household income (%)** |  |  |  | < .001 |
| Less than \$10,000 | 17 (6.2%) | 15 (18%) | 2 (1.0%) | |
| \$10,000–24,999 | 17 (6.2%) | 14 (17%) | 3 (1.6%) | |
| \$25,000–49,999 | 27 (9.9%) | 14 (17%) | 13 (6.8%) | |
| \$50,000–74,999 | 27 (9.9%) | 15 (18%) | 12 (6.3%) | |
| \$75,000–99,999 | 29 (11%) | 7 (8.4%) | 22 (12%) | |
| \$100,000–199,999 | 93 (34%) | 12 (14%) | 81 (42%) | |
| \$200,000 or more | 64 (23%) | 6 (7.2%) | 58 (30%) | |

*Note.* Values are n (%) for categorical variables and M (SD) for continuous variables. *p*-values are from Pearson's chi-squared test, Fisher's exact test, or Wilcoxon rank-sum test as appropriate.

### Measures

#### Paid family leave

At the V05 study visit, caregivers retrospectively reported whether they took leave from work following the birth of their child and, if so, whether that leave was paid, unpaid, or a combination of both. Consistent with prior work (e.g., Brito et al., 2022; Kozak et al., 2021), caregivers who reported paid leave or a combination of paid and unpaid leave were classified as having PFL (n = 191); caregivers who reported no leave or unpaid leave only were classified as not having PFL (n = 83). An analogous set of items assessed whether the infant’s father or other co-parent took paid leave and the highest level of education they had completed.

#### Household demographics and covariates

Maternal education was assessed at the V01 visit as the highest level of education completed, coded on a 0–21-point scale spanning no formal schooling to a doctoral degree (*M* = 17.7, *SD* = 2.3, approximately a 4-year college degree). Father or partner education was assessed at V05 using the same scale. Household income was assessed at V01 as one of seven ordinal brackets ranging from less than $10,000 to $200,000 or more. Gestational age at delivery (in weeks) and child sex were obtained from the V02 visit. Prenatal substance exposure, used only in the sensitivity analyses reported below, was operationalized as a binary composite indicating any positive screen or caregiver-reported use of nicotine, opioids, stimulants, alcohol, or cannabis, drawn from clinical and timeline follow-back measures collected at V01.

#### Language outcome

Language ability was assessed at the V04 visit (M adjusted age = 12.2 months, SD = 1.4) using the Language Standard Score of the Bayley Scales of Infant and Toddler Development, Fourth Edition (Bayley-4; Bayley & Aylward, 2019), a standardized, normed (M = 100, SD = 15) measure of receptive and expressive communication administered by trained examiners.

#### Diffusion MRI and fractional anisotropy

Diffusion-weighted images were acquired at the V03 visit (M adjusted age = 5.9 months, SD = 1.5) using the HBCD Study’s harmonized, multi-vendor diffusion MRI protocol, modeled closely on the Adolescent Brain Cognitive Development (ABCD) Study protocol (Dean et al., 2024). The protocol acquires 1.7-mm isotropic contiguous axial slices (87 slices; 238 x 238 mm field of view; TR = 4800 ms; TE = 88 ms; multiband acceleration factor 3) using multi-shell diffusion encoding at b = 0, 500, 1000, 2000, and 3000 s/mm², split across two series with reversed phase encoding for a combined total of 130 diffusion-encoding directions, for a total acquisition time of approximately 12.5 minutes (Dean et al., 2024). Diffusion data were preprocessed using QSIPrep’s infant-specific processing workflow (Dean et al., 2024). Fractional anisotropy (FA), a scalar diffusion MRI metric reflecting the directional coherence of water diffusion within white matter and commonly used as an index of white matter microstructural organization, was extracted along the left arcuate fasciculus, left inferior longitudinal fasciculus, and left uncinate fasciculus from the HBCD Study’s standardized diffusion processing outputs. Analyses focused on the left hemisphere given its established role in language lateralization (Zuk et al., 2021).

### Analytic Approach

All analyses were conducted in version 4.5.3 of R. All dependent variables were visually inspected for normality using histograms, and any outliers exceeding 3 standard deviations from the mean were winsorized. For each outcome (Bayley-4 Language Standard Score; FA in the arcuate fasciculus, inferior longitudinal fasciculus, and uncinate fasciculus), three nested linear mixed-effects models were estimated with study site as a random intercept, using maximum-likelihood estimation (lme4, lmerTest packages in R): Model 1 included covariates only. Model 2 added the main effect of PFL. Model 3 added the interaction between PFL and maternal education. Follow-up simple-slope contrasts (emmeans, interactions packages) estimated the association between PFL and each outcome at maternal education levels one standard deviation below the mean, at the mean, and one standard deviation above the mean. Tables report the main-effects model (Model 2) and the interaction model (Model 3) separately for each outcome.

In parallel, a second set of models substituted paternal or partner PFL as the predictor and paternal or partner education as the moderator (and covariate, in place of maternal education), estimated in the subsample with complete paternal/partner leave data, using the same covariate set and model sequence.

The covariate set was specified a priori to mirror the set used by Kozak et al. (2021): gestational age at delivery, child sex, the corresponding parent’s education, and household income. For the FA outcomes, adjusted age at the V03 MRI visit was added as a covariate, because FA increases steeply with age across this interval. No age-at-assessment covariate was included in the Bayley-4 models, because the Language Standard Score is already normed for age. A random intercept for site was added because the HBCD Study recruits across 27 sites that differ in scanner hardware, recruitment catchment, and state paid-leave policy.

As a sensitivity analysis, we refit all models containing a PFL term with prenatal substance-use exposure as an additional covariate. HBCD intentionally recruits families with prenatal substance exposure, making it useful to assess whether adjustment changed the PFL estimates.

## Results

Table 1 presents sample characteristics by maternal PFL status. Table 2 presents descriptive statistics for study outcomes among the Bayley-4 (*n* = 231) and diffusion MRI (*n* = 148) subsamples. Groups did not differ significantly at the bivariate level on any outcome measure (all ps > .05).

**Table 2.** Outcome Variables by Maternal Paid Family Leave (PFL) Status.

| **Outcome** | **Overall** | **No Paid Leave** | **Paid Leave** | ***p*** |
| --- | --- | --- | --- | --- |
| **Bayley-4 subsample (n = 231)** |  |  |  |  |
| Adjusted age at V04, months, M (SD) | 12.2 (1.4) | 12.0 (1.6) | 12.3 (1.3) | .2 |
| Bayley-4 Language Standard Score, M (SD) | 99.1 (11.1) | 99.5 (11.7) | 98.9 (10.9) | .5 |
| **Diffusion MRI (FA) subsample (n = 148)** |  |  |  |  |
| Adjusted age at V03 MRI, months, M (SD) | 5.9 (1.5) | 5.7 (1.3) | 6.0 (1.6) | .5 |
| FA, left arcuate fasciculus, M (SD) | 0.222 (0.018) | 0.218 (0.018) | 0.224 (0.020) | .062 |
| FA, left inferior longitudinal fasciculus, M (SD) | 0.215 (0.014) | 0.212 (0.014) | 0.216 (0.015) | .2 |
| FA, left uncinate fasciculus, M (SD) | 0.194 (0.012) | 0.192 (0.012) | 0.194 (0.011) | .4 |

*Note.* FA = fractional anisotropy. The Bayley-4 and diffusion MRI subsamples are non-overlapping outcome subsamples of the primary analytic sample (N = 274) and are presented separately. p-values are from Pearson's chi-squared test, Fisher's exact test, or Wilcoxon rank-sum test as appropriate.

### Maternal Paid Leave, Language, and White Matter

#### Bayley-4 Language Standard Score

Maternal PFL was not associated with language ability at approximately 12 months. The main effect of PFL was nonsignificant (Model 2: *b* = – 1.201, *SE* = 1.843, *t*(226.0) = –0.65, *p* = .516; Table 3), as was its interaction with maternal education (Model 3: *b* = 0.729, *SE* = 0.701, *t*(227.3) = 1.04, *p* = .299; Table 4). Simple-slope contrasts gave no evidence of an association at any level of maternal education (–1 SD, 15.4 years: *Δ* = –2.71, *SE* = 2.34, *t* = –1.16, *p* = .25; mean, 17.7 years: *Δ* = –1.01, *SE* = 1.85, *t* = –0.55, *p* = .58; +1 SD, 20.1 years: *Δ* = 0.69, *SE* = 2.58, *t* = 0.27, *p* = .79).

**Table 3.**
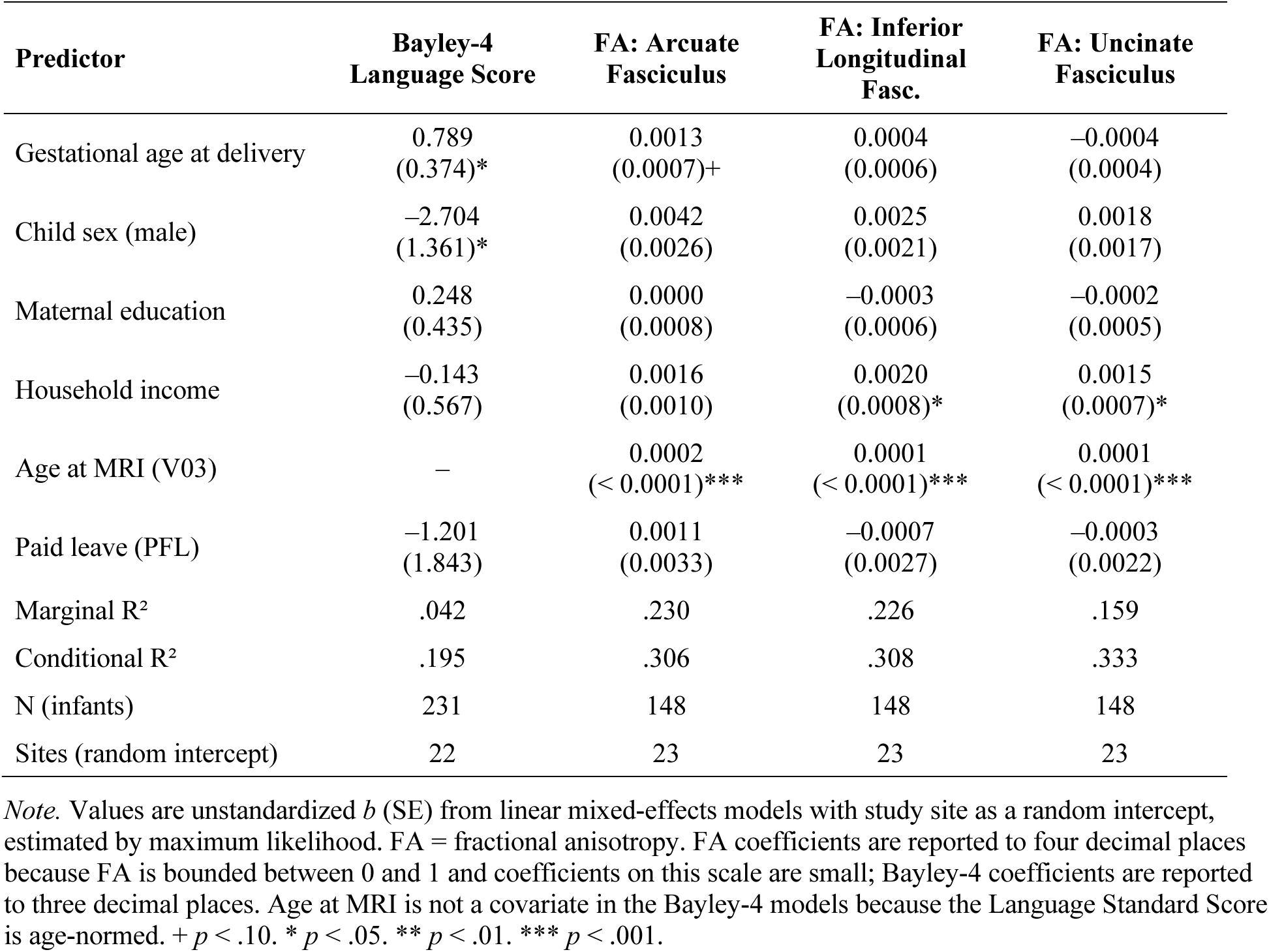
Linear Mixed-Effects Models (Model 2: Covariates + Main Effect of Maternal PFL) Predicting Language and White Matter Outcomes.

**Table 4.**
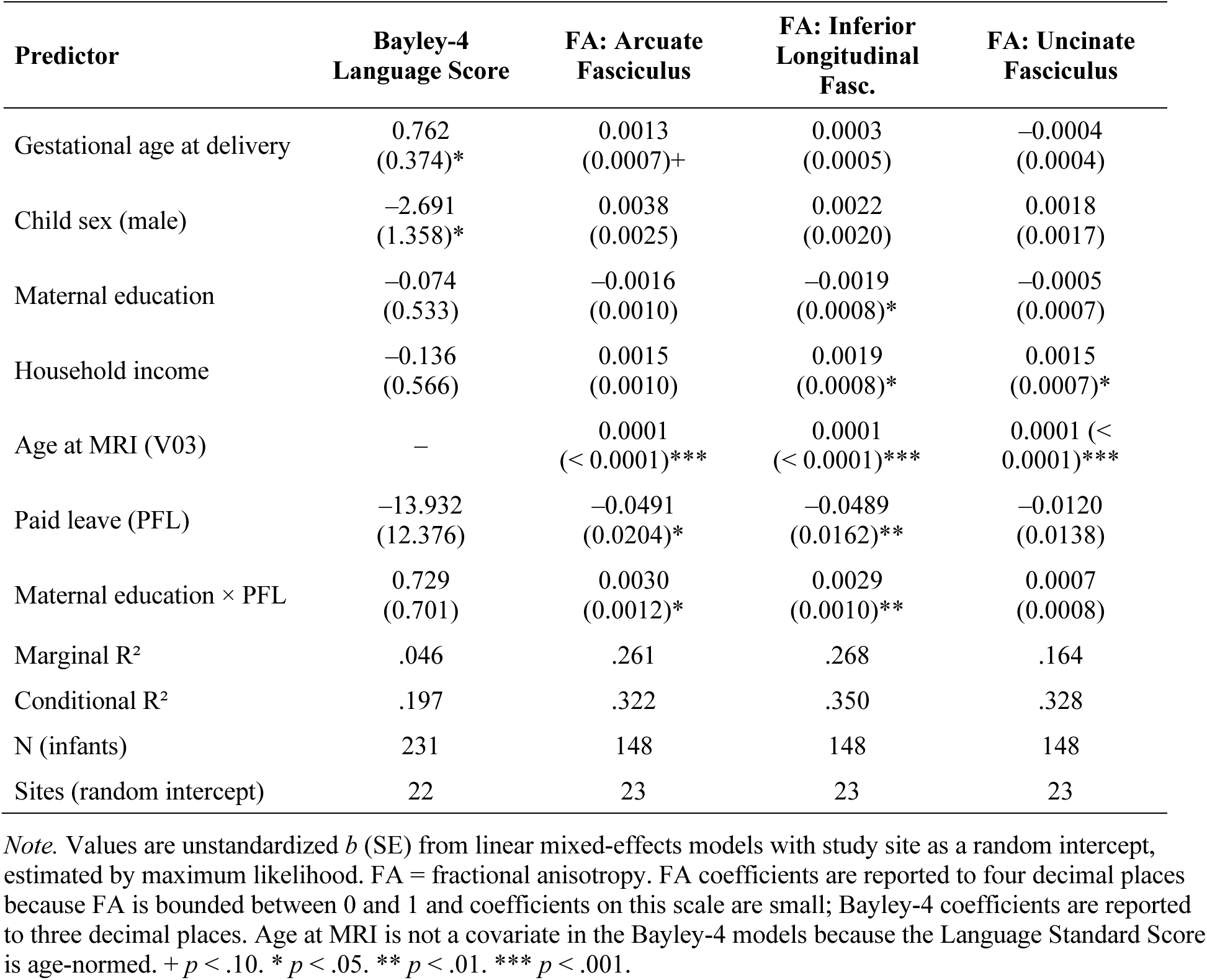
Linear Mixed-Effects Models (Model 3: Covariates + Maternal PFL × Maternal Education) Predicting Language and White Matter Outcomes.

#### Fractional anisotropy: Arcuate fasciculus

In the main-effects model, PFL was not independently associated with arcuate fasciculus FA (Model 2: *b* = 0.0011, *SE* = 0.0033, *t*(147.0) = 0.32, *p* = .751; Table 3). The interaction between PFL and maternal education was significant (Model 3: *b* = 0.0030, *SE* = 0.0012, *t*(145.2) = 2.49, *p* = .014; Table 4), such that the association between PFL and FA became more positive as maternal education increased. Simple-slope contrasts indicated that PFL was associated with significantly higher FA among infants of mothers with relatively high educational attainment (+1 SD, 20.0 years: *t* = 2.09, *p* = .04), but not among infants of mothers with average (17.5 years: *t* = 0.89, *p* = .37) or relatively low (–1 SD, 15.0 years: *t* = –1.15, *p* = .25) educational attainment (Figure 1).

**Figure 1.**
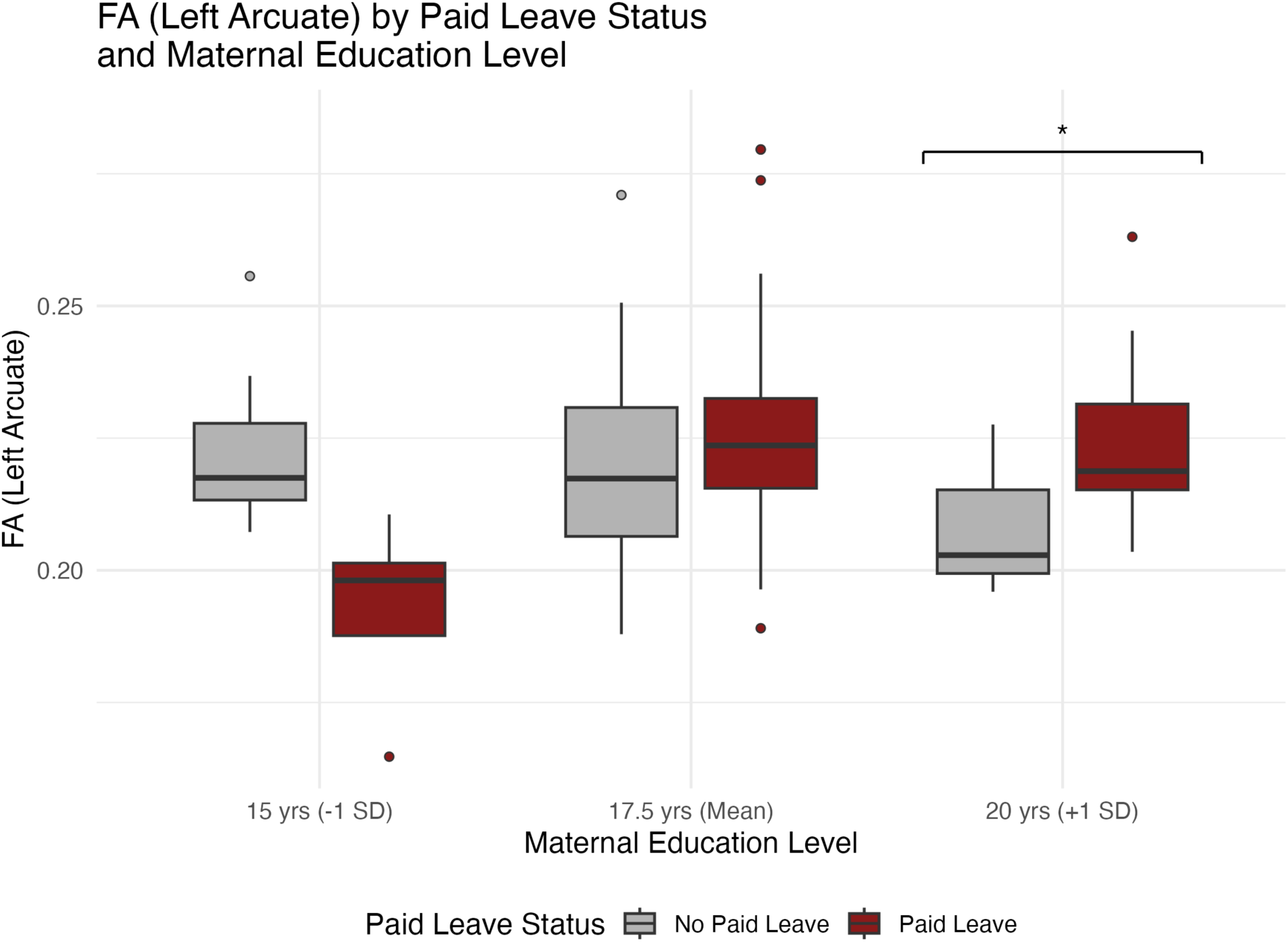
Predicted fractional anisotropy (FA) in the left arcuate fasciculus by maternal paid family leave (PFL) status and maternal education level, based on estimated marginal means from Model 3 (Table 4). Error bars represent 95% confidence intervals; the asterisk indicates a significant paid-leave contrast (*p* = .04) at +1 SD maternal education.

#### Fractional anisotropy: Inferior longitudinal fasciculus

A similar pattern emerged for the inferior longitudinal fasciculus. PFL was not independently associated with FA in the main-effects model (Model 2: *b* = –0.0007, *SE* = 0.0027, *t*(147.1) = –0.27, *p* = .789; Table 3), but the PFL × maternal education interaction was significant (Model 3: *b* = 0.0029, *SE* = 0.0010, *t*(146.7) = 3.00, *p* = .003; Table 4). Simple-slope contrasts indicated that PFL was associated with significantly higher FA among infants of mothers with relatively high educational attainment (+1 SD: *t* = 2.11, *p* = .04), showed no association at average maternal education (*t* = 0.45, *p* = .65), and showed a nonsignificant trend in the opposite direction at low maternal education (–1 SD: *t* = –1.92, *p* = .06; Figure 2).

**Figure 2.**
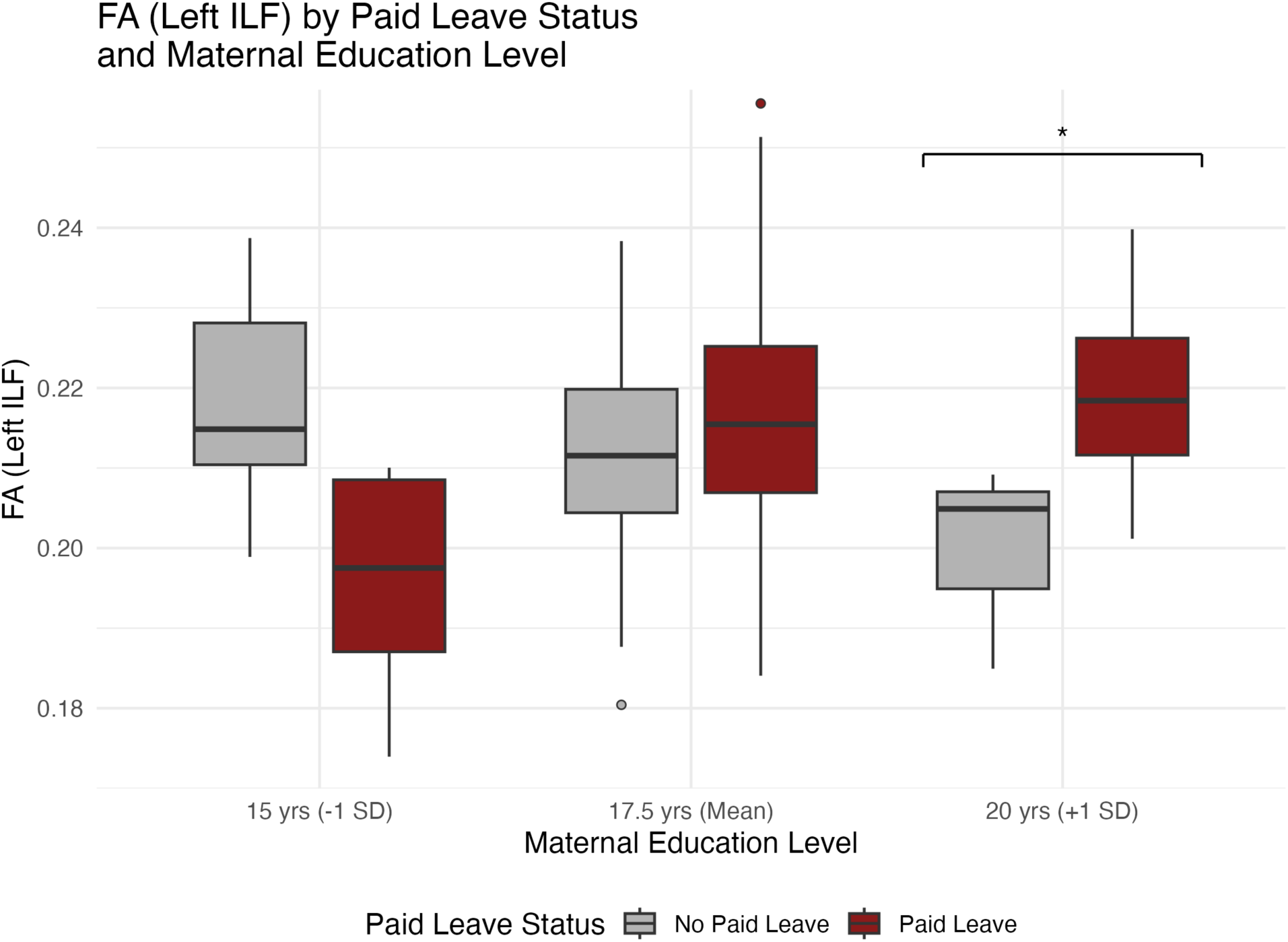
Predicted fractional anisotropy (FA) in the left inferior longitudinal fasciculus by maternal paid family leave (PFL) status and maternal education level, based on estimated marginal means from Model 3 (Table 4). Error bars represent 95% confidence intervals; the asterisk indicates a significant paid-leave contrast (*p* = .04) at +1 SD maternal education.

#### Fractional anisotropy: Uncinate fasciculus

Contrary to our hypothesis, neither the main effect of PFL (Model 2: *b* = –0.0003, *SE* = 0.0022, *t*(147.9) = –0.16, *p* = .873; Table 3) nor its interaction with maternal education (Model 3: *b* = 0.0007, *SE* = 0.0008, *t*(147.8) = 0.85, *p* = .394; Table 4) was associated with FA in the uncinate fasciculus, and no simple-slope contrast approached significance (*ps* ≥ .53).

### Paternal Paid Leave, Language, and White Matter

Parallel models substituting paternal PFL as the predictor and paternal education as the moderator produced no significant main effect of leave for any outcome in the main-effects models (Supplemental Table 1; all *ps* ≥ .311). In the interaction models, moderation by father or partner education was nonsignificant for the Bayley-4 Language Standard Score (Model 3: *b* = 0.679, *SE* = 0.741, *t*(186.7) = 0.92, *p* = .361), the arcuate fasciculus (*b* = 0.0002, *SE* = 0.0014, *t*(107.5) = 0.17, *p* = .869), and the inferior longitudinal fasciculus (*b* = 0.0001, *SE* = 0.0011, *t*(104.8) = 0.13, *p* = .896; Supplemental Table 2).

For the uncinate fasciculus, the paternal PFL × education interaction reached the conventional threshold, (*b* = –0.0017, *SE* = 0.0008, *t*(107.2) = –1.99, *p* = .049). No simple-slope contrast was significant at any level of father or partner education, however (–1 SD, 14.9 years: *t* = 0.80, *p* = .42; mean, 17.4 years: *t* = –0.76, *p* = .45; +1 SD, 19.8 years: *t* = –1.59, *p* = .11). We therefore treat this interaction as a trend rather than an interpretable moderation effect.

### Sensitivity Analyses: Adjusting for Prenatal Substance-Use Exposure

Every model containing a PFL term was refit with prenatal substance-use exposure added to the a priori covariate set. Substance-exposure data were complete in both outcome subsamples, so these models were fit on the same samples as the primary models (*n* = 231 for the Bayley-4 and *n* = 148 for FA in the maternal models; *n* = 193 and *n* = 120 in the paternal or partner models). Full coefficients appear in Supplemental Tables 3 and 4.

Adding prenatal substance-use exposure left every PFL estimate essentially unchanged. In the maternal models, the PFL × maternal education interaction remained significant and nearly identical in magnitude for the arcuate fasciculus (*b* = 0.0030, *SE* = 0.0012, *t*(145.1) = 2.49, *p* = .014) and the inferior longitudinal fasciculus (*b* = 0.0029, *SE* = 0.0010, *t*(146.7) = 3.00, *p* = .003), and remained nonsignificant for the uncinate fasciculus (*b* = 0.0007, *SE* = 0.0008, *p* = .395) and the Bayley-4 Language Standard Score (*b* = 0.734, *SE* = 0.690, *p* = .288). Simple-slope patterns were likewise unchanged, with a significant paid-leave contrast only at high maternal education for the arcuate (*p* = .04) and inferior longitudinal (*p* = .03) fasciculi. In the paternal or partner models, the uncinate interaction remained at the conventional threshold (*b* = –0.0018, *SE* = 0.0008, *t*(107.3) = –2.09, *p* = .039) with no significant simple slopes (*ps* ≥ .09), and all other paternal or partner estimates remained nonsignificant.

## Discussion

In a national, multi-site U.S. sample, maternal PFL was not directly associated with infant language outcomes or white matter microstructure. However, maternal education moderated associations between PFL and FA in two of the three hypothesized language-relevant white matter tracts: paid leave was associated with higher FA in the left arcuate fasciculus and inferior longitudinal fasciculus only among infants whose mothers had relatively high educational attainment. Contrary to our hypothesis, no comparable moderation was observed for the uncinate fasciculus, or for the Bayley-4 Language Standard Score. Parallel analyses testing paternal PFL, to our knowledge the first test of this question for any brain outcome, did not show a comparable pattern for any tract, aside from a marginal, opposite-direction trend for the uncinate fasciculus. These findings extend prior evidence linking PFL to infant brain function (Brito et al., 2022) and toddler language outcomes (Kozak et al., 2021) by identifying a specific, tract-level structural correlate of maternal PFL, showing that this correlate is not distributed equally across families, and providing a first, if inconclusive, look at whether paternal or partner leave-taking relates to the same outcomes.

That paid leave showed no direct association with language at 12 months deserves emphasis, because language is the outcome for which prior evidence is strongest. Kozak et al. (2021) reported higher parent-reported language scores among toddlers whose mothers took paid leave, and we did not reproduce that association using an examiner-administered standard score at approximately 12 months of age. Several differences could account for this. Their outcome was assessed later in development and by parent report, whereas ours was assessed at an age when expressive language is only beginning to emerge and when the Bayley-4 Language Standard Score consequently has limited predictive utility to later outcomes (Rubio-Cordina & Grantham-McGregor, 2020). It is also possible that structural differences detectable at 6 months precede detectable differences in behavioral language ability, which typically becomes more differentiated, and more reliably measured, later in the second year of life. Prior work linking early white matter organization to later, rather than concurrent, language outcomes (Zuk et al., 2021) suggests that longer-term follow-up within HBCD may be needed to determine whether the structural differences identified here translate into detectable behavioral differences as this cohort ages.

The pattern of results is broadly consistent with a resource-amplification account, in which families with greater educational and, likely, associated economic and social resources are better positioned to convert the time afforded by PFL into caregiving experiences that support white matter development. These may include more consistent routines, greater access to high-quality caregiving resources and educational materials in the home, and/or lower financial strain during the leave period itself. This pattern diverges from Kozak et al.’s (2021) finding that PFL was most beneficial for toddler socioemotional outcomes among lower-income families, and from Brito et al.’s (2022) finding of no moderation by family income for infant EEG outcomes. Together with the present results, this heterogeneity across studies suggests that the extent to which, and the families for whom, PFL confers benefit may depend on the developmental outcome under study, the specific indicator of family resources considered, and possibly on the structure of the PFL policies available to a given sample. Rather than resolving this heterogeneity, the present findings underscore that PFL should not be assumed to operate as a uniformly protective exposure, and that policy efforts to expand access to PFL may need to be paired with complementary supports if the goal is to narrow, rather than widen, existing disparities in early child development.

We had hypothesized that PFL would relate to FA in all three tracts, including the uncinate fasciculus, given its documented role in semantic association and lexical retrieval in addition to its more prominent roles in episodic memory and socioemotional processing (Papagno, 2011; Von Der Heide et al., 2013). That this hypothesis was not supported may reflect genuine differences in how these tracts relate to early caregiving experience, or it may reflect measurement factors specific to the uncinate fasciculus. Relative to the arcuate fasciculus and inferior longitudinal fasciculus, the uncinate fasciculus follows a particularly protracted developmental course, continuing to mature well beyond infancy and into adulthood (Lebel & Beaulieu, 2011), and its hook-like anatomy has long made it comparatively difficult to reconstruct reliably with tractography (Von Der Heide et al., 2013). Both factors could reduce our ability to detect an association that, if present, may not yet be developmentally expressed, or reliably measurable, at 6 months of age.

This study’s parallel test of paternal or partner PFL, motivated by the near-total absence of research on non-maternal caregivers’ leave-taking and infant brain development despite evidence that paternal leave and involvement relate to child development more broadly (Huerta et al., 2013; Kim et al., 2016), did not reproduce the pattern observed for maternal leave. No significant associations emerged for language or for the arcuate or inferior longitudinal fasciculus. The one exception was the uncinate fasciculus, where the interaction between paternal or partner leave and father or partner education reached *p* = .049 in the direction opposite to the maternal pattern. We do not treat this as a finding. No simple-slope contrast was significant at any level of father or partner education, which is the form the effect would need to take to be substantively meaningful. Given that fathers and partners in this sample took paid leave under different policy and workplace conditions than mothers, and that the subsample with complete father/partner data was smaller (*n* = 120–193) and possibly systematically different from the full sample, these null and marginal results should not be interpreted as evidence that paternal or partner caregiving is unimportant for infant brain development. Rather, they represent, to our knowledge, a first direct test of this question and underscore the need for larger, purpose-designed studies of paternal and non-maternal caregiver leave, which remains understudied relative to maternal leave in both the economics and developmental literatures.

Several limitations qualify these findings. First, PFL was assessed via retrospective caregiver report at a visit occurring after the leave period, which may be subject to recall error, and did not capture the source of wage replacement (e.g., employer policy, state paid-leave program, or short-term disability insurance), information that may be relevant given substantial state-level variation in PFL policy. Second, as in nearly all observational studies of PFL, families who take paid leave differ systematically from those who do not on multiple, likely only partially measured, dimensions (Table 2); although models adjusted for gestational age at delivery, child sex, maternal education, and household income, and although adding prenatal substance-use exposure did not alter any conclusion, residual confounding by unmeasured family or workplace characteristics cannot be ruled out, and causal interpretation of these associations is not warranted. Third, this study tested a relatively large number of tracts, outcomes, and interaction terms without formal preregistration or correction for multiple comparisons; although the covariate set was fixed a priori, these analyses are exploratory, and the specific p-values reported here should be interpreted as hypothesis-generating rather than confirmatory, pending replication in an independent sample or in the fuller HBCD cohort as data collection continues. Fourth, sample sizes for imaging (*n* = 148) and father/partner analyses (*n* = 120–193) were modest for detecting interaction effects, which typically require larger samples than main effects to estimate reliably.

This study drew on a subsample of the HBCD Study cohort, restricted to families with complete leave, covariate, and outcome data. Relative to the broader U.S. birth population, this subsample was disproportionately college-educated and White, likely reflecting both patterns of retention in a demanding longitudinal neuroimaging study and the sociodemographic patterning of PFL access itself. Because families with the least access to PFL, and arguably the most to gain from it, may be systematically underrepresented in analyses that require complete leave and neuroimaging data, the present findings may understate the true heterogeneity of PFL’s association with infant brain development across the full range of U.S. families. As HBCD Study data collection continues and the sample grows, particularly at sites serving more socioeconomically and racially/ethnically diverse communities, larger and more representative subsamples will allow for a fuller test of whether, and for whom, PFL relates to infant brain and language development.

Paid family leave is increasingly discussed as a lever for reducing early childhood disparities. The present findings suggest that its association with infant white matter development, at least in this sample, was concentrated among already-advantaged families, a pattern that, if replicated, would caution against treating expanded access to PFL alone as sufficient to narrow early developmental disparities. Understanding which features of PFL, duration, wage replacement, job protection, or the caregiving practices it enables, drive benefit for which families will be essential for designing policy that supports equitable early brain development.

## Open Practices Statement

The data necessary to reproduce the analyses presented here are not publicly accessible in the specific extracted form used for this manuscript; de-identified HBCD Study data are available to qualified researchers through the NIMH Data Archive following the consortium’s data use certification process (Nelson et al., 2024). The analytic code (R Markdown) necessary to reproduce the analyses presented here is available from the corresponding author upon request. The materials necessary to attempt to replicate the findings presented here are not publicly accessible beyond the HBCD Study’s published protocol documentation (docs.hbcdstudy.org). The analyses presented here were not preregistered.

---

**Supplemental Table 1.** Secondary Linear Mixed-Effects Models (Model 2: Covariates + Main Effect of Paternal/Partner PFL) Predicting Language and White Matter Outcomes.

| Predictor | Bayley-4<br>Language Score | FA: Arcuate<br>Fasciculus | FA: Inferior<br>Longitudinal<br>Fasc. | FA: Uncinate<br>Fasciculus |
| --- | --- | --- | --- | --- |
| Gestational age at delivery | 0.885<br>(0.389)* | 0.0010<br>(0.0007) | 0.0004<br>(0.0006) | −0.0008<br>(0.0005)+ |
| Child sex (male) | −2.424<br>(1.447)+ | 0.0036<br>(0.0028) | 0.0029<br>(0.0023) | 0.0021<br>(0.0018) |
| Paternal education | −0.133<br>(0.400) | −0.0002<br>(0.0007) | 0.0001<br>(0.0006) | 0.0001<br>(0.0005) |
| Household income | −0.881<br>(0.647) | 0.0023<br>(0.0013)+ | 0.0013<br>(0.0010) | 0.0018<br>(0.0008)* |
| Age at MRI (V03) | — | 0.0002<br>($< 0.0001$***) | 0.0001<br>($< 0.0001$***) | 0.0001<br>($< 0.0001$***) |
| Paternal paid leave (PFL) | 2.027<br>(1.996) | 0.0005<br>(0.0039) | 0.0004<br>(0.0031) | −0.0002<br>(0.0025) |
| Marginal R<sup>2</sup> | .053 | .282 | .242 | .225 |
| Conditional R<sup>2</sup> | .187 | .355 | .378 | .393 |
| N (infants) | 193 | 120 | 120 | 120 |
| Sites (random intercept) | 20 | 20 | 20 | 20 |

*Note.* Values are unstandardized $b$ (SE) from linear mixed-effects models with study site as a random intercept, estimated by maximum likelihood, in the secondary subsample with complete father/partner leave data. FA = fractional anisotropy. FA coefficients are reported to four decimal places because FA is bounded between 0 and 1 and coefficients on this scale are small; Bayley-4 coefficients are reported to three decimal places. + $p < .10$ . \* $p < .05$ . \*\* $p < .01$ . \*\*\* $p < .001$ .

**Supplemental Table 2.**
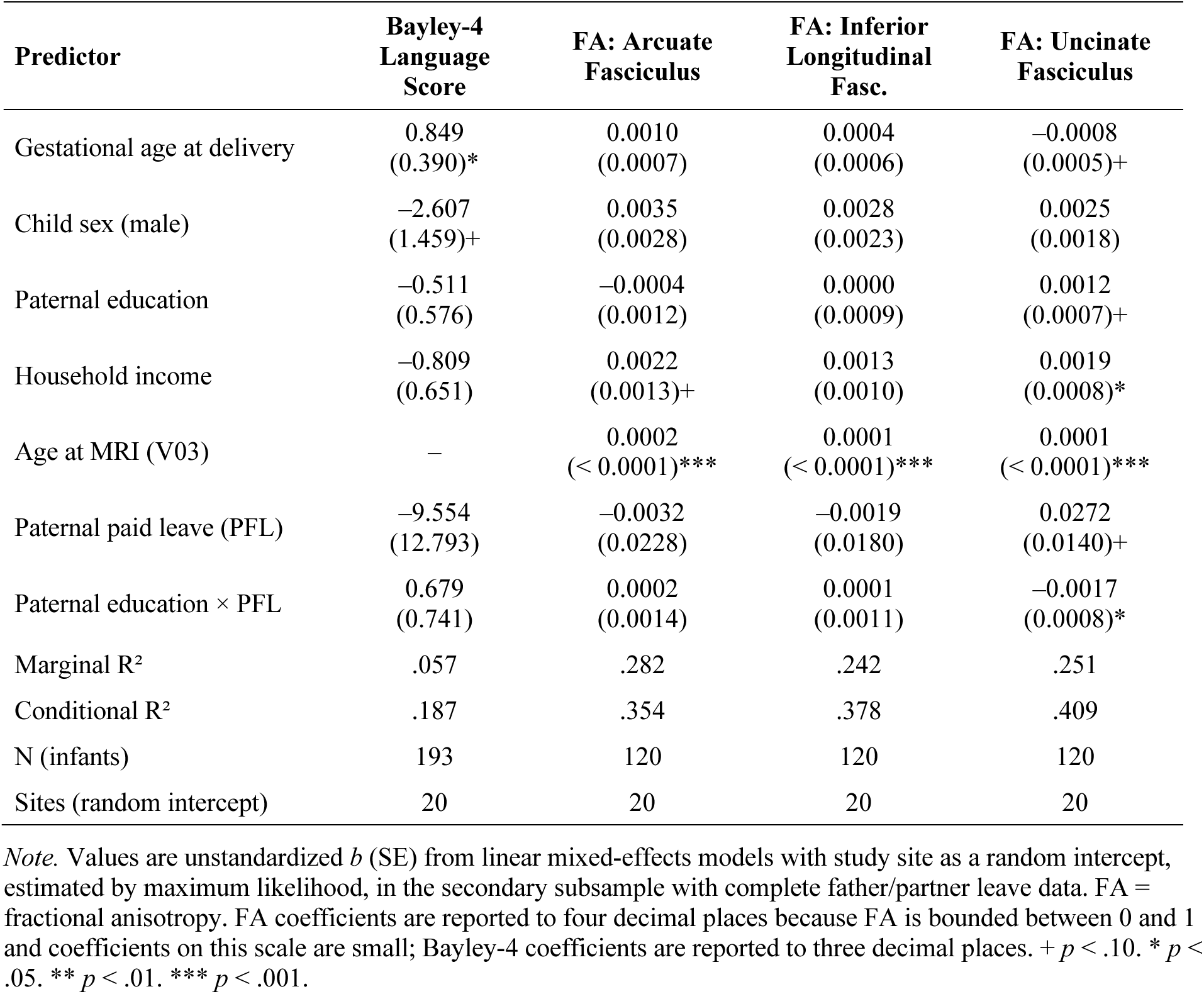
Secondary Linear Mixed-Effects Models (Model 3: Covariates + Paternal/Partner PFL × Father/Partner Education) Predicting Language and White Matter Outcomes.

**Supplemental Table 3.** Sensitivity Analyses: Maternal PFL Models Adjusting Additionally for Prenatal Substance-Use Exposure.

| Predictor | Bayley-4<br>Language Score | FA: Arcuate<br>Fasciculus | FA: Inferior<br>Longitudinal<br>Fasc. | FA: Uncinate<br>Fasciculus |
| --- | --- | --- | --- | --- |
| Gestational age at delivery | 0.671<br>(0.370)+ | 0.0012<br>(0.0007)+ | 0.0003<br>(0.0005) | −0.0004<br>(0.0005) |
| Child sex (male) | −2.651<br>(1.337)* | 0.0039<br>(0.0025) | 0.0022<br>(0.0020) | 0.0018<br>(0.0017) |
| Maternal education | 0.115<br>(0.529) | −0.0016<br>(0.0010) | −0.0019<br>(0.0008)* | −0.0006<br>(0.0007) |
| Household income | −0.005<br>(0.559) | 0.0015<br>(0.0010) | 0.0019<br>(0.0008)* | 0.0014<br>(0.0007)* |
| Age at MRI (V03) | — | 0.0001<br>($< 0.0001$)*** | 0.0001<br>($< 0.0001$)*** | 0.0001<br>($< 0.0001$)*** |
| Prenatal substance-use<br>exposure | 4.673<br>(1.665)** | 0.0011<br>(0.0030) | 0.0010<br>(0.0024) | −0.0007<br>(0.0020) |
| Paid leave (PFL) | −13.694<br>(12.178) | −0.0489<br>(0.0204)* | −0.0487<br>(0.0162)** | −0.0120<br>(0.0138) |
| Maternal education × PFL | 0.734<br>(0.690) | 0.0030<br>(0.0012)* | 0.0029<br>(0.0010)** | 0.0007<br>(0.0008) |
| Marginal R<sup>2</sup> | .075 | .262 | .269 | .164 |
| Conditional R<sup>2</sup> | .216 | .323 | .351 | .330 |
| N (infants) | 231 | 148 | 148 | 148 |
| Sites (random intercept) | 22 | 23 | 23 | 23 |

**Supplemental Table 4.**
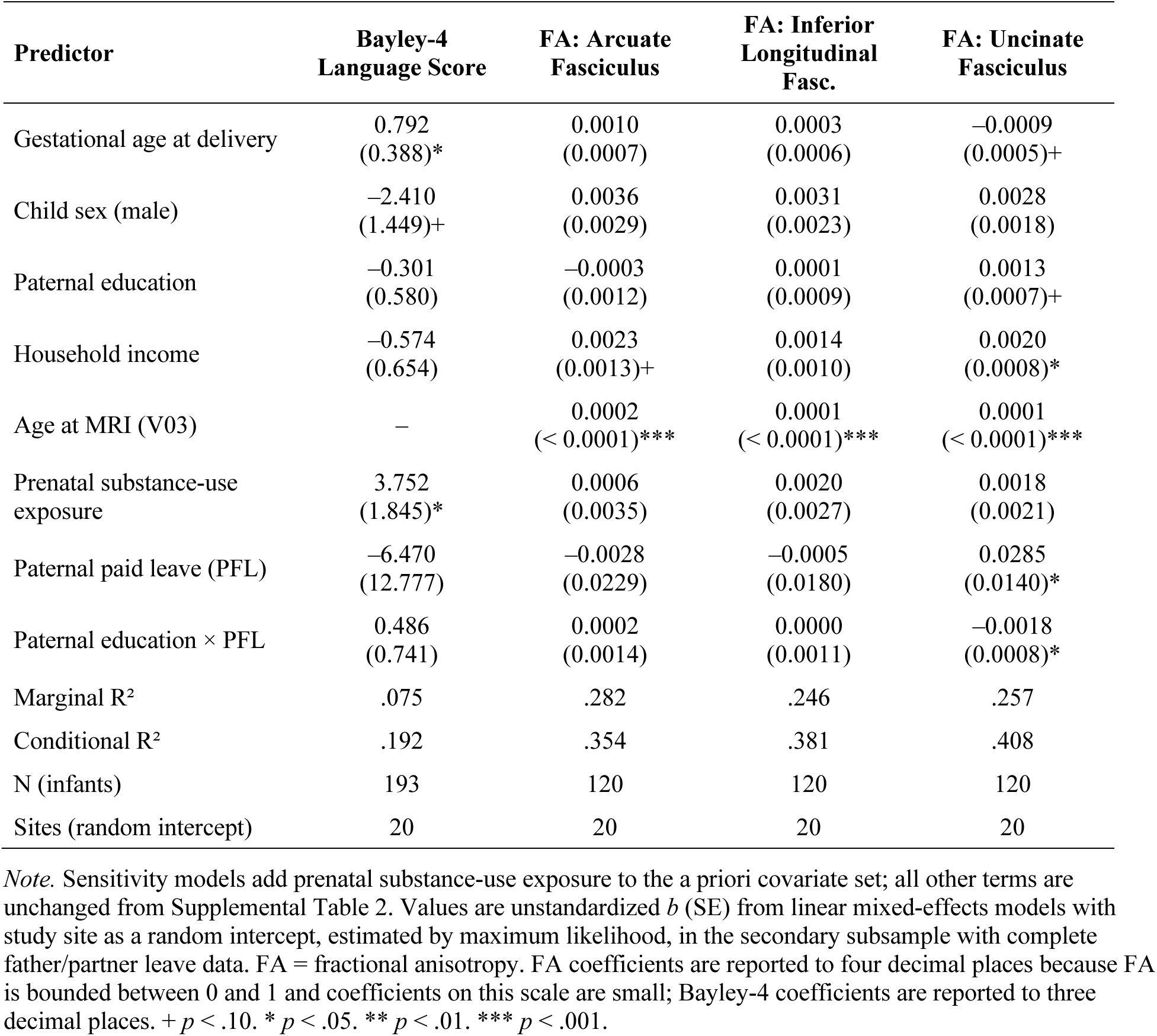
Sensitivity Analyses: Paternal/Partner PFL Models Adjusting Additionally for Prenatal Substance-Use Exposure.

| Predictor | Bayley-4<br>Language Score | FA: Arcuate<br>Fasciculus | FA: Inferior<br>Longitudinal<br>Fasc. | FA: Uncinate<br>Fasciculus |
| --- | --- | --- | --- | --- |
| Gestational age at delivery | 0.792<br>(0.388)* | 0.0010<br>(0.0007) | 0.0003<br>(0.0006) | −0.0009<br>(0.0005)+ |
| Child sex (male) | −2.410<br>(1.449)+ | 0.0036<br>(0.0029) | 0.0031<br>(0.0023) | 0.0028<br>(0.0018) |
| Paternal education | −0.301<br>(0.580) | −0.0003<br>(0.0012) | 0.0001<br>(0.0009) | 0.0013<br>(0.0007)+ |
| Household income | −0.574<br>(0.654) | 0.0023<br>(0.0013)+ | 0.0014<br>(0.0010) | 0.0020<br>(0.0008)* |
| Age at MRI (V03) | — | 0.0002<br>($< 0.0001$)*** | 0.0001<br>($< 0.0001$)*** | 0.0001<br>($< 0.0001$)*** |
| Prenatal substance-use<br>exposure | 3.752<br>(1.845)* | 0.0006<br>(0.0035) | 0.0020<br>(0.0027) | 0.0018<br>(0.0021) |
| Paternal paid leave (PFL) | −6.470<br>(12.777) | −0.0028<br>(0.0229) | −0.0005<br>(0.0180) | 0.0285<br>(0.0140)* |
| Paternal education × PFL | 0.486<br>(0.741) | 0.0002<br>(0.0014) | 0.0000<br>(0.0011) | −0.0018<br>(0.0008)* |
| Marginal R<sup>2</sup> | .075 | .282 | .246 | .257 |
| Conditional R<sup>2</sup> | .192 | .354 | .381 | .408 |
| N (infants) | 193 | 120 | 120 | 120 |
| Sites (random intercept) | 20 | 20 | 20 | 20 |

